# Adaptive cytoskeletal responses to extracellular environment viscosity modulate cell migration

**DOI:** 10.1101/2025.07.01.662417

**Authors:** Zhongya Lin, Xindong Chen, Xi-Qiao Feng

**Affiliations:** Institute of Biomechanics and Medical Engineering, AML, Department of Engineering Mechanics, Tsinghua University, Beijing 100084, China; Mechano-X Institute, Tsinghua University, Beijing 100084, China

**Keywords:** viscosity, actin filament dynamics, cell migration, short-term memory

## Abstract

Cell migration is a pivotal process in metastasis, allowing cancer cells to invade surrounding tissues and disseminate to distant organs. While extracellular environment (ECE) viscosity serves as a critical modulator of cell motility, its regulatory mechanisms remain unclear. This study presents a mechanobiological model to investigate how ECE viscosity modulates cancer cell migration by regulating some key processes, including actin polymerization, retrograde flow, and adhesion adaptations. Our results reveal a biphasic response: a moderate increase in ECE viscosity enhances actin filament network density and adhesion strength, thereby accelerating migration, whereas excessively high viscosity hinders movement due to too large mechanical resistance. Furthermore, we identify a short-term migration memory phenomenon, where cancer cells exposed to high viscosity environments retain elevated migration speeds after transitioning to low viscosity conditions. This memory effect is sustained by the continued assembly of cytoskeletal proteins such as actin monomers and Arp2/3. These analyses reveal an adaptive mechano-chemo-biological mechanism by which cancer cells integrate and respond to mechanical cues from their viscous environment to optimize migration, and advance the understanding of cancer cell migration in various tissue environments.

## Introduction

Cell migration underpins numerous physiological and pathological phenomena. Examples include the migration of stem cells to establish the body plan and organ systems during the earliest stages of embryogenesis^1,2^, the intricate movements of immune cells in response to infection^3,4^, and the complex dynamics of cancer metastasis^5,6^. Actin cytoskeleton is a primary force-generating structure that drives cell motility, mainly through dynamic assembly and disassembly processes at the leading edge of migrating cells^7–9^. Actin filaments (AFs) rapidly polymerize to form structures such as lamellipodia and filopodia, which are essential for protrusion and adhesion to the extracellular environment^10,11^. Cells explore their environment and adapt their subsequent responses through dynamic cytoskeletal processes. Therefore, the physical properties of the extracellular environment, including stiffness, topography, crosslinking, viscoelasticity, and porosity, significantly influence cellular behavior and actin dynamics^12–17^. Among these factors, the role of viscosity has emerged as a critical but relatively understudied modulator of cell motility, actively shaping the biomechanical behavior of cells^18–20^.

Elevated viscosity has been observed in various types of tumors, often due to the accumulation of cancer cells and their secreted factors^16,21^. Interestingly, recent studies have shown that increased ECE viscosity can enhance the motility of certain cancer cell types^20,22,23^, challenging the conventional expectation that higher resistance would suppress cell movement. This counterintuitive finding suggests that cancer cells have evolved adaptive mechanisms to navigate more efficiently through viscous environments, which may be driven by the alterations in actin dynamics, ion channel activity, and cell-matrix interactions. Furthermore, the changes in ECE viscosity may lead to migration memory, where cancer cells retain their migratory behavior after exposure to different viscosities^23^. Mechanical memory may significantly enhance their ability to survive and thrive in distant organs, whose mechanisms remain to be elucidated^24–26^. Therefore, a thorough understanding of how ECE viscosity modulates cell migration is crucial for decoding the mechanisms that drive tumor progression and metastasis.

Although the importance of actin dynamics in cell motility has been well recognized, the effects of varying ECE viscosity on the organization and behavior of the AF network have not been thoroughly investigated. Furthermore, while several recent works have addressed how viscosity influences actin dynamics, they overlooked the critical feedback mechanisms between cellular components that lead to mechanical memory. To address this gap, we establish a multiscale mechanobiological model that simulates the evolution of the AF network at the leading edge in response to changes in ECE viscosity. We aim to elucidate how variations in viscosity influence actin polymerization, retrograde flow, and cell-matrix adhesions, thereby characterizing the mechanistic influences of ECE viscosity on cancer cell migration. Furthermore, we investigate how exposure to different viscosities can induce lasting biomechanical adaptations, i.e., cytoskeleton-based short-term migration memory, in cancer cells, providing new insights into their ability to optimize movement in response to different mechanical cues.

## Results

### Mechanobiological model of cell migration

In this paper, we focus on lamellipodia induced migration. Lamellipodia are characterized by a branched network of AFs, primarily formed through the action of Arp2/3, which serves as a nucleation site for new filament assembly^27,28^. This protein promotes the formation of a dense network that drives the protrusion of the cell membrane. For a cancer cell migrating in the viscous ECE, the driving force is provided by the branched AF network near the leading edge, which generates sufficient propulsion for cell movement (Fig. 1a). The leading edge is subject to the propulsive force of AF polymerization, the resistance of cell membrane tension and the ECE (Fig. 1b). At the steady state, the force balance equation at the leading edge of a migrating cell is

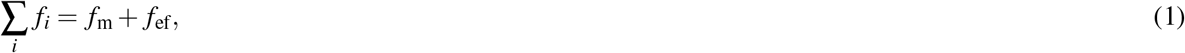

where *f*_*i*_ is the propulsive force of the *i*th AF. *f*_m_ and *f*_ef_ denote the membrane tension and the resistance of the viscous ECE, respectively. Here, the membrane tension is assumed to be constant during cell migration, and we have also discussed the effect of its variation with ECE viscosity on cell migration (Fig. S5). The resistance from the viscous ECE can be written as a function of viscosity *µ* and migration speed *v*

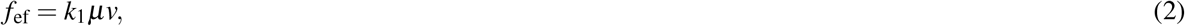

where *k*_1_ is a dimensionless scaling constant.

**Fig. 1.**
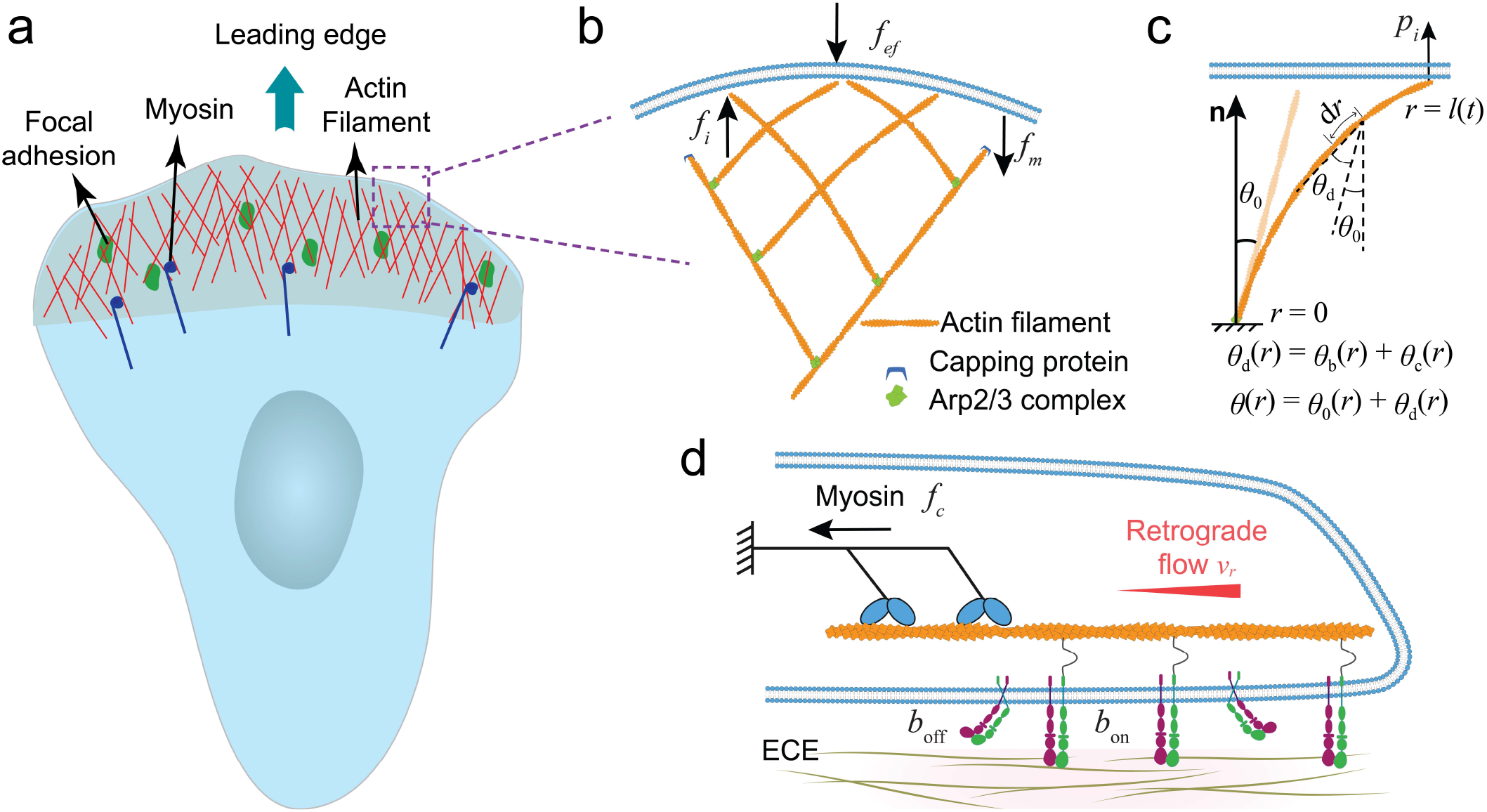
Schematic representation of AF dynamics during cell migration. (a) Formation of the AF network at the leading edge. (b) Localized view of the branched AF network. The AF deforms as it interacts with the cell membrane, and the deformation influences its subsequent branching behavior. (c) Deformation of an AF and its interaction with the cell membrane. The AF is considered as a beam that can withstand bending and shear deformation. (d) Molecular clutch model with viscous ECE. Myosin contractility pulls on the AFs, leading to retrograde flow, and integrins bound to the ECE provide resistance. AF: actin filament.

The polymerization process of AFs is characterized by the addition of individual actin monomers to the barbed ends of an AF. This process is regulated by biological, chemical and mechanical cues^7,29^. Accordingly, the growth rate of the polymerizing AF is expressed as^30,31^

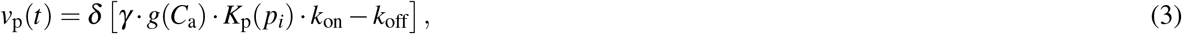

where *δ* is the diameter of an actin monomer, *C*_a_ = *C*_a_(Φ) is the local concentration of actin monomers around the leading edge, which is proportional to the polymerized AF density Φ, *g*(*C*_a_) denotes the polymerization rate is a function of actin monomer concentration, *γ* is the diffusion coefficient of actin monomers, *k*_on_ and *k*_off_ are the association and dissociation rates of actin monomers to the polymerizing AF, respectively, *K*_p_ is the force-dependent probability of AF polymerization. Thus, the length of the polymerizing AF is

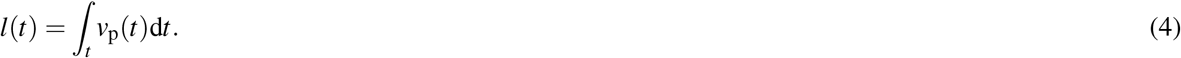

Previous studies indicated that the load-induced curvature of AFs can influence the formation of the branched AF network at the leading edge^32,33^. Recent cryo-EM experiments demonstrated that the mother filament in contact with the Arp2/3 complex exhibits slight bending and twisting^34^, as shown in Fig. S1. In this study, the AF is treated as a deformable beam with a circular cross-section. When it interacts with the leading edge membrane, it undergoes bending and shear deformation (Fig. 1c). Thus, the interaction force acting on the *i*th AF is

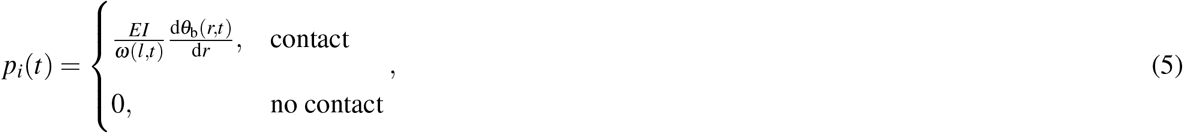

where *E* is the Young’s modulus of the AF, *I* = *πd*^4^*/*64 is the second moment of the cross-sectional area of the AF with diameter *d, θ*_b_ is the bending angle, and *ω*(*l, t*) is the deflection. The propulsive force is *f*_*i*_(*t*) = *p*_*i*_(*t*) cos φ, where φ denotes the angle between the normal direction of the local leading edge and the direction of cell migration. In addition, the bending force induces conformational changes of the AF, affecting its branching behavior^35^. The new branch prefers the convex side of a bent filament over the concave side^30,33^. That is, the Arp2/3 complex, as an actin nucleator, has a higher binding affinity to the convex side of an AF. A bending curvature dependent factor *s*_arp_(*κ*_m_) is introduced in Eq. 7, which denotes the distance between two adjacent Arp2/3 complex branches along a mother AF, where *κ*_m_ is the mean bending curvature of the AF.

Moreover, we consider the retrograde flow that occurs in the AFs within the lamellipodia of migrating cells, mainly due to myosin contraction^36,37^. The pulling force of myosin motors on the AF, driving the actin flow toward the cell center (Figs. 1d and S2). Integrins, which link the actin cytoskeleton to the extracellular environment and provide effective mechanotransduction^38^, serve as molecular clutches. The retrograde flow speed can be described by the Hill relation^11,39^:

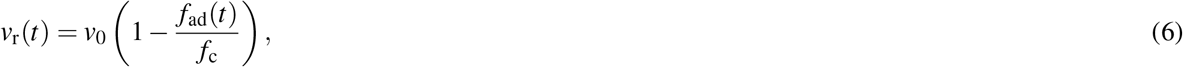

where *v*_0_ is the free flow speed, *f*_c_ is the myosin stalling force, and *f*_ad_ is the adhesion force generated by the engaged clutches. Besides, force transmission can lead to strengthening and stabilization of adhesions^40^. In our model, adhesion strengthening is captured by the addition of new clutches when a threshold force acting on the binding clutches is reached (Eq. S10, and Fig. S2). Thus, our model integrates the effects of AF polymerization, branching and retrograde flow, which occur in cell migration. The simulation procedure is described in the Methods section and Fig. S11. The detailed derivation of this mechanobiological model can be found in the Supporting Information.

We first calculate the cytoskeleton polymerization rate under different viscous environments, corresponding to different resistances, and compare the obtained resistance-growth rate relation with relevant experiments (Figs. S3 and S4). The results show that our model can capture the force-sensitive polymerization behavior, i.e., a slower growth rate under higher resistance conditions, which is consistent with the experimental results^26^. Our model has accounted for adhesion and actin retrograde flow, which regulate cell motility with cytoskeleton dynamics together.

### Increasing ECE viscosity leads to biphasic migration speed

To investigate the effects of increasing ECE viscosity on cancer cell migration, we employ our mechanobiological model to simulate the response of migration speed, focal adhesion dynamics, and retrograde actin flow under varying viscosity conditions. The simulations reveal a biphasic response in the migration speed as ECE viscosity increases (Figs. 2a and S5.). Specifically, the migration speed initially increases, reaching a peak around 700 cP, before decreasing at much higher viscosity. This suggests that moderate increase in viscosity enhances cell motility, which is consistent with experimental observations^22^. However, beyond this optimal viscosity range, excessive resistance impedes movement, akin to the well-established effects of resistance on cell motility^41^. Our model further shows that the adaptive response of a cell is closely linked to changes in its cytoskeletal organization. As viscosity increases, the higher resistance prompts an increase in the number of AFs at the leading edge (Figs. 2b and S3). The increase in filament density and corresponding forces suggests that the cell compensates for the elevated mechanical resistance by recruiting more cytoskeletal elements to maintain sufficient propulsive forces for forward movement. Meanwhile, in a high viscosity environment (e.g., 370 cP), cells exhibit larger focal adhesions (engaged clutches) compared to those in a low viscosity environment (1 cP) (Fig. 2c, left). This indicates an adaptive response wherein cells strengthen their adhesion in high viscosity conditions to sustain propulsive forces during migration. Experimental data on different cells supports this trend, demonstrating an increased number of focal adhesions per cell in high viscosity conditions (Fig. 2c, right;^18^).

**Fig. 2.**
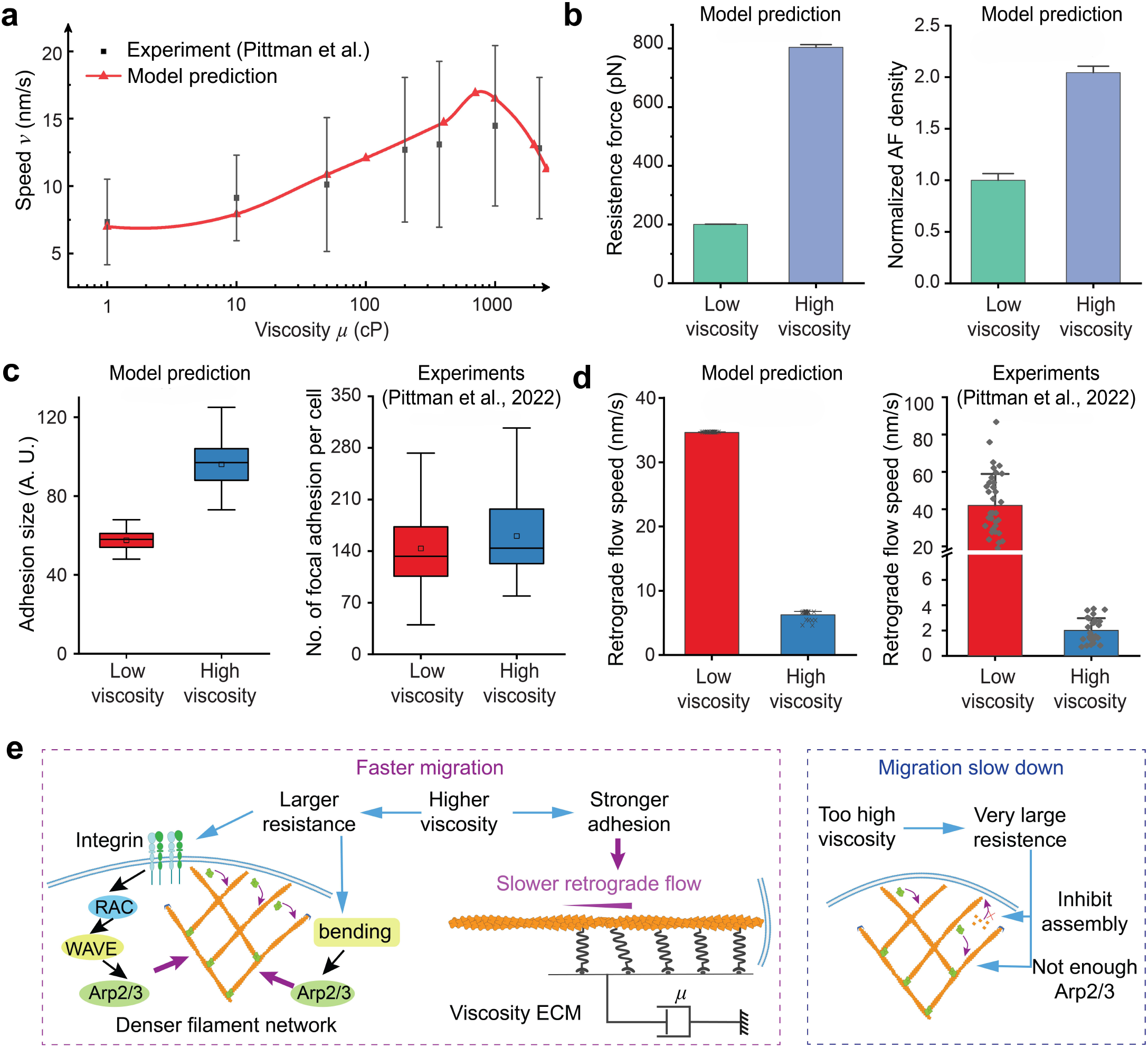
Differences in the process of cell migration caused by ECE viscosity. (a) Increasing and then decreasing migration speed with increasing ECE viscosity. The simulation results correspond to the average migration speed. The experimental results are the motility of MDA-MB-231 breast cancer cells in viscous media from Pittman et al.^22^. (b) The model prediction of the resistance force (left) and the AF number (right) at the leading edge. The AF density denotes the number of AFs whose point end is no more than 200nm from the leading edge. (c) Viscosity induced focal adhesion adaptation. The model prediction indicates that the larger adhesion size in the high viscosity medium (370 cP) compared to the low viscosity medium (1 cP). The adhesion size corresponds to the number of engaged clutches during the simulation. The experiment corresponds to the number of focal adhesions per cell in the low viscosity case (regular medium, 1 cP) and high viscosity case (added 1% high-molecular-weight hydroxypropyl methylcellulose, 370 cP). (d) Effect of viscosity on retrograde flow. The model prediction indicates that the retrograde flow is slower in the high viscosity medium (370 cP) compared to the less viscous medium (1 cP). The experiment corresponds to the same tendency, that is, the retrograde flow is much slower in the high viscosity case (370 cP) than in the low viscosity case (1 cP). (e) Mechanobiological mechanism of ECE viscosity on cell motility. The migration speed corresponds to the statistics after the simulation is stabilized.

Additionally, our simulations reveal a marked decrease in retrograde flow speed with increasing viscosity, consistent with experimental observations (Fig. 2d). Retrograde flow is significantly slower in high viscosity media (370 cP) compared to low viscosity media (1 cP). This reduction in retrograde flow speed corresponds to a mechanical adaptation of the actin cytoskeleton to heightened adhesions, reflecting a shift in the extent to which the cytoskeleton transmits forces under different conditions. Therefore, the mechanobiological feedback mechanism can be summarized (Fig. 2e). In moderately viscous ECE, the increased resistance enhances cell migration by promoting larger adhesions, denser AF network, and reduced retrograde flow. However, in environments of excessive viscosity, migration speed is ultimately constrained by overwhelming mechanical resistance, which inhibits actin polymerization and diminishes the efficiency of force transmission necessary for cell movement.

### Deformation of AFs influences the network density

Each AF is modeled as a bending beam to explore its dynamic responses to different ECE viscosities. We analyze the changes in AF curvature, density, and the resulting propulsive forces under different viscosity conditions. Simulations reveal that AFs exhibit greater curvatures in high viscosity environments compared to those in low viscosity conditions (Fig. 3a). This increased curvature is a direct consequence of the increased mechanical resistance imposed by the viscous medium, which forces the AFs to bend further. The bending of the AFs facilitates a higher probability of Arp2/3 complex binding to the convex side of the curved AFs, as governed by Eq. (7). As a result, the number of bound Arp2/3 complexes, responsible for nucleation of new AFs at the leading edge, is significantly higher in high viscosity environments (Fig. 3b). This contributes to a denser AF network, which is necessary to generate the propulsive forces required for membrane protrusion under higher resistance conditions. Our results show a strong positive correlation between the curvature of AF and the AF density at the leading edge (Fig. 3c), where increasing AF curvature leads to a significant increase in AF density. This indicates that mechanical bending of AFs directly increases AF density, enabling the cell to adapt to the elevated mechanical resistance.

**Fig. 3.**
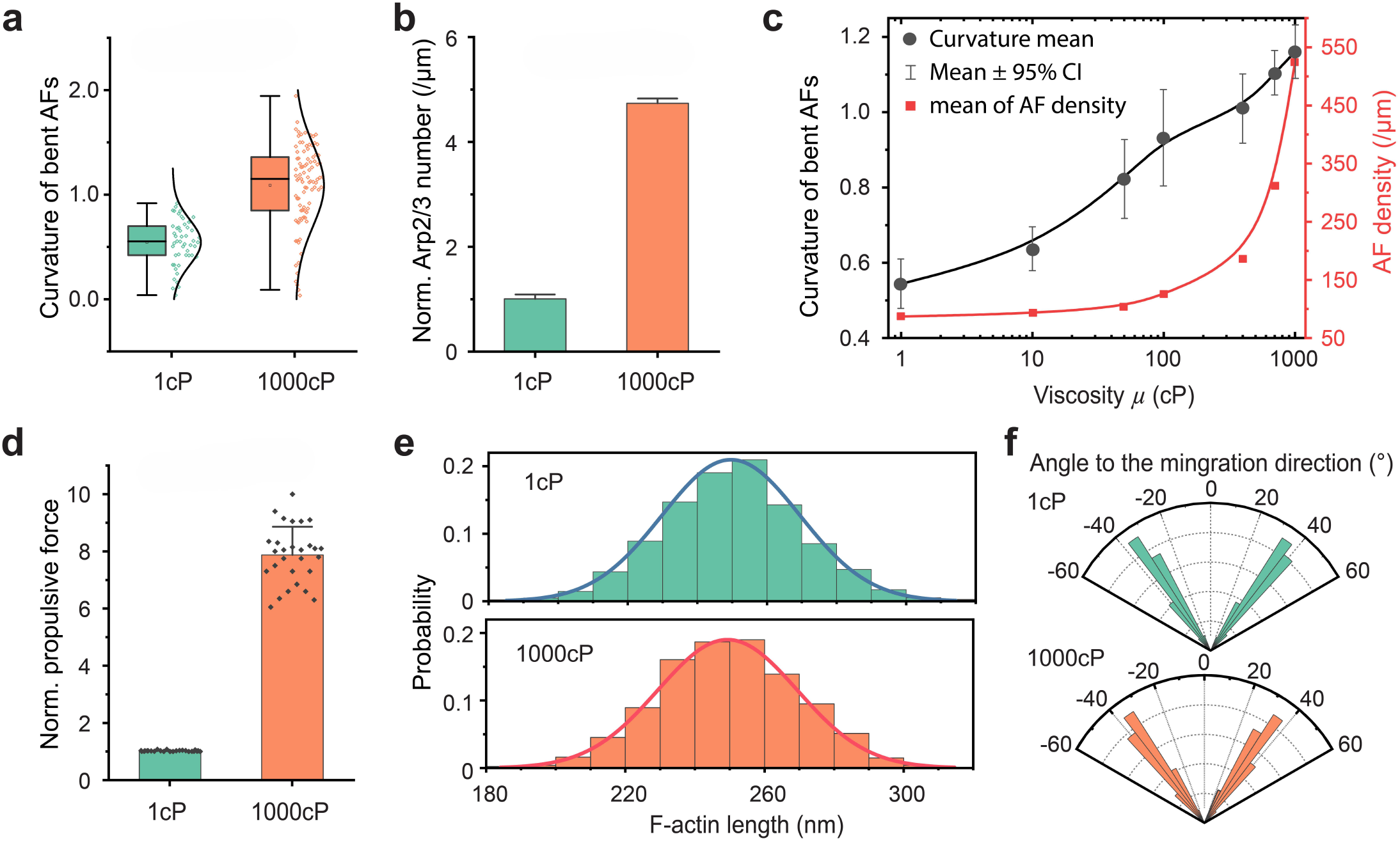
Deformation and distribution of AFs. (a) The mean curvature of bent AFs in low and high viscosity ECE. The curvature is much larger in the latter condition. (b) The normalized number of Arp2/3 bound to the mother AFs. More bound Arp2/3 is related to more newly generated daughter AFs, that is, a denser AF network. Error bars, s.d. (c) The relation between the curvature of the bent AF and its density at the leading edge. The AF density is proportional to the statistical mean curvature. (d) The propulsive force attributed by bent AFs. The denser AF network and larger curvature in the high viscosity ECE lead to a larger propulsive force. Error bars, s.d. (e) The length of an individual AF follows a Gaussian distribution, and (f) at an approximate angle of ±35^°^ from the forward direction of migration.

Moreover, the propulsive force generated by the bent AFs is significantly greater in the high viscosity conditions (Fig. 3d). The combination of increased AF curvature and a denser filament network produces a larger force to drive membrane protrusion, a critical adaptation to overcome the mechanical challenges of high viscosity environments. The lengths of individual AFs follow a Gaussian distribution in all conditions (Fig. 3e), and the initial direction before bending is aligned within an angular range of approximately ±35° from the forward migration direction (Fig. 3f). Additionally, filament length also influences cellular and molecular behaviors, with longer filaments supporting increased cell migration and contributing to a denser AF network (Fig. S6). These results highlight the critical interplay between filament geometry and network density in regulating the cellular response to mechanical resistance. Such insights offer a deeper understanding of how cancer cells adapt their cytoskeletal structure to optimize migration in challenging microenvironments.

### Actin monomer concentration influences migration speed

The availability of profilin-bound actin monomers significantly affects AF polymerization. Our simulations show that as ECE viscosity increases, the actin monomer concentration near the leading edge is upregulated to sustain AF polymerization (Fig. 4a). Within a given viscosity condition, the concentration of actin monomers affects AF polymerization and then acts as a change in migration speed (Fig. 4b). Increased availability of actin monomers enhances the polymerization of AFs, which provides propulsive forces necessary for efficient movement, thus leading to faster migration speeds. To further elucidate the role of actin monomer availability, we examined three specific cases (Fig. 4c). Normally, the monomer concentration is below 22.5µM under low viscosity conditions (1 cP and 100 cP), as shown in Fig. 4a. Increasing the monomer concentration to 22.5µM represents an elevated supply. While in the high viscosity condition (1000 cP), the monomer concentration normally exceeds 22.5µM. Reducing the available monomer concentration to 22.5µM in this setting limits the availability of actin monomer. Fig. 4d, illustrating the migration speeds, shows that an increased supply of actin monomers significantly boosts the migration speed, whereas a reduced availability of actin monomers significantly slows the migration.

**Fig. 4.**
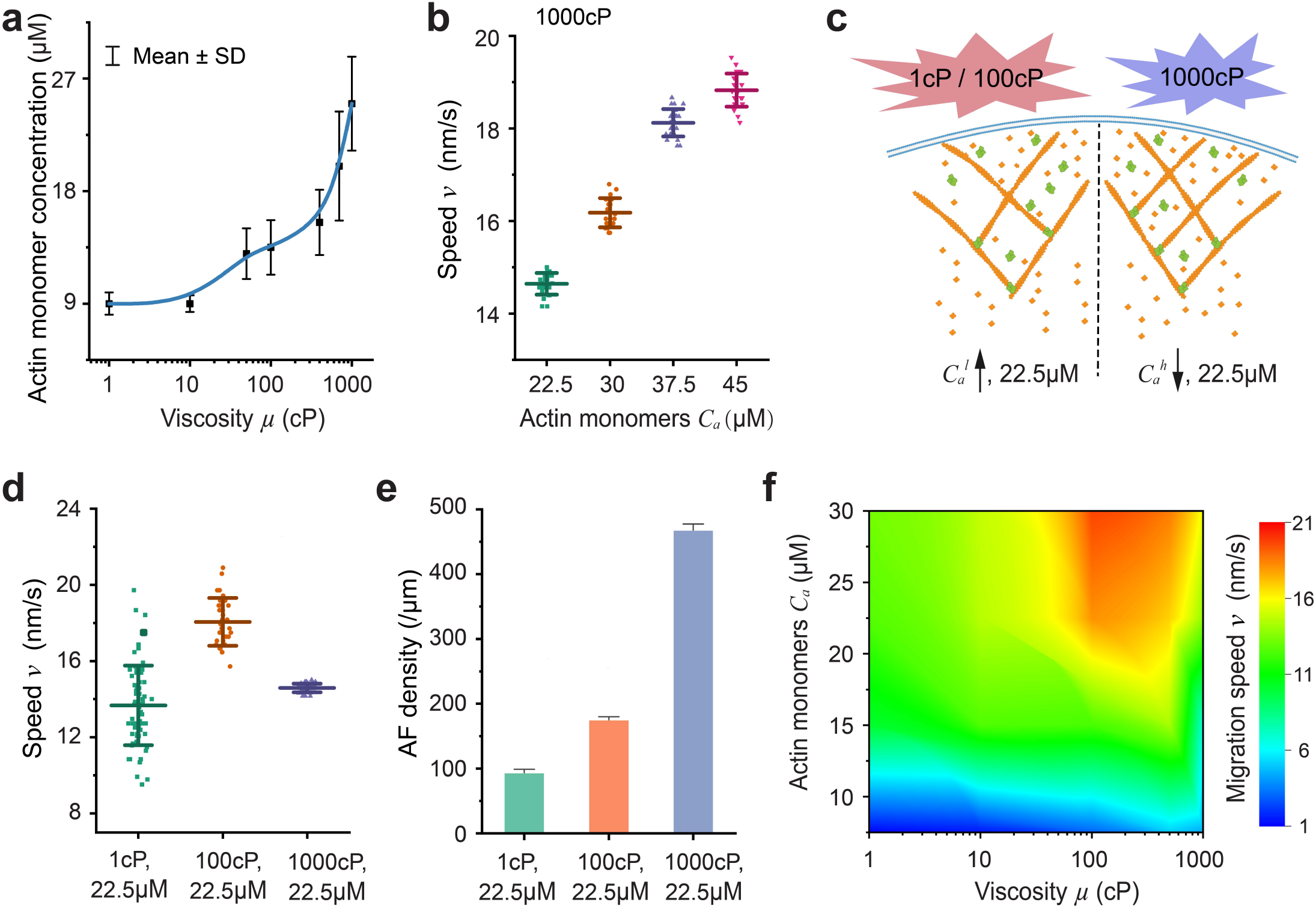
Influence of actin monomers and viscosity on the AF network and cell migration. (a) Actin monomer concentration at each step for different viscosity environments. Error bars, s.d. (b) Influence of actin monomer concentration on migration speed. Higher concentration promotes AF polymerization, leading to higher migration speed. (c) Cases simulating the influence of actin monomers on cell behavior. For lower viscosity cases (1 cP/100 cP), the actin monomer concentration is less than 22.5*µ*M in normal situation, thus, the concentration 22.5*µ*M means increasing the supply of actin monomers. In contrast, the concentration 22.5*µ*M means reducing the supply of actin monomers for the high viscosity case (1000 cP). (d) Migration speed of the three cases presented in (c). (e) The AF density at the leading edge of the three cases. (f) Phase diagram of biochemical cues for cell migration speed, which shows the optimal ECE viscosity and actin monomer concentration for cell migration.

The corresponding AF density at the leading edge (Fig. 4e) shows that the AF density is higher in the low viscosity conditions (1 cP and 100 cP) with elevated actin monomer supply (22.5µM) compared to the normal condition (Fig. 3c). In contrast, reducing the monomer supply to 22.5µM in the high viscosity condition (1000 cP) results in a lower AF density compared to the normal condition. These AF adjustments correspond to changes in cell motility. Then, we present a phase diagram that maps the interplay between ECE viscosity and actin monomer availability, delineating the optimal conditions for maximizing cell migration speed (Fig. 4f). It shows that low monomer concentrations or excessively high viscosity can impair movement, while an optimal combination of these factors supports rapid migration. These results emphasize that effective cell migration depends on a tuned balance between the availability of associated proteins and the mechanical resistance provided by the extracellular environment, offering insights into how cancer cells adapt their motility to varying physical conditions.

### Integrin ligand density and myosin contraction influence retrograde flow

Integrin ligand density and myosin concentration play critical roles in modulating retrograde flow and cell motility through mechanobiological interactions such as cell-ECE adhesion, integrin activation, and myosin-induced contractility^42–44^. Our results show that cell-ECE adhesion becomes stronger with increasing ECE viscosity (Fig. 5a, left). This increased adhesion contributes to a reduction in retrograde flow speed under higher viscosity conditions (Fig. 5a, right). This slowed retrograde flow enables the generation of more effective propulsive forces to drive forward movement. We further investigate how ligand density, which refers to the availability of binding sites such as integrins or adhesion complexes^45^, influences retrograde flow speed and migration speed. When ligand density is low, retrograde flow speed is higher, indicating weaker cell-ECE adhesion and allowing the actin network to move backwards more rapidly (Fig. 5b, left). This condition corresponds to the reduced migration speed (5b, right). However, as the ligand density increases, the retrograde flow decreases, and the cell migration speed increases due to the enhanced propulsion of AF networks. Interestingly, beyond a certain threshold, further increases in ligand density have little effect on migration speed (5b, right), indicating a plateau where adhesion strength is maximized and additional binding sites do not significantly enhance motility. This phenomenon is in partial agreement with experimental results^46^, while the biphasic dependence of cell migration speed on ligand density^46,47^ requires further modeling investigation.

**Fig. 5.**
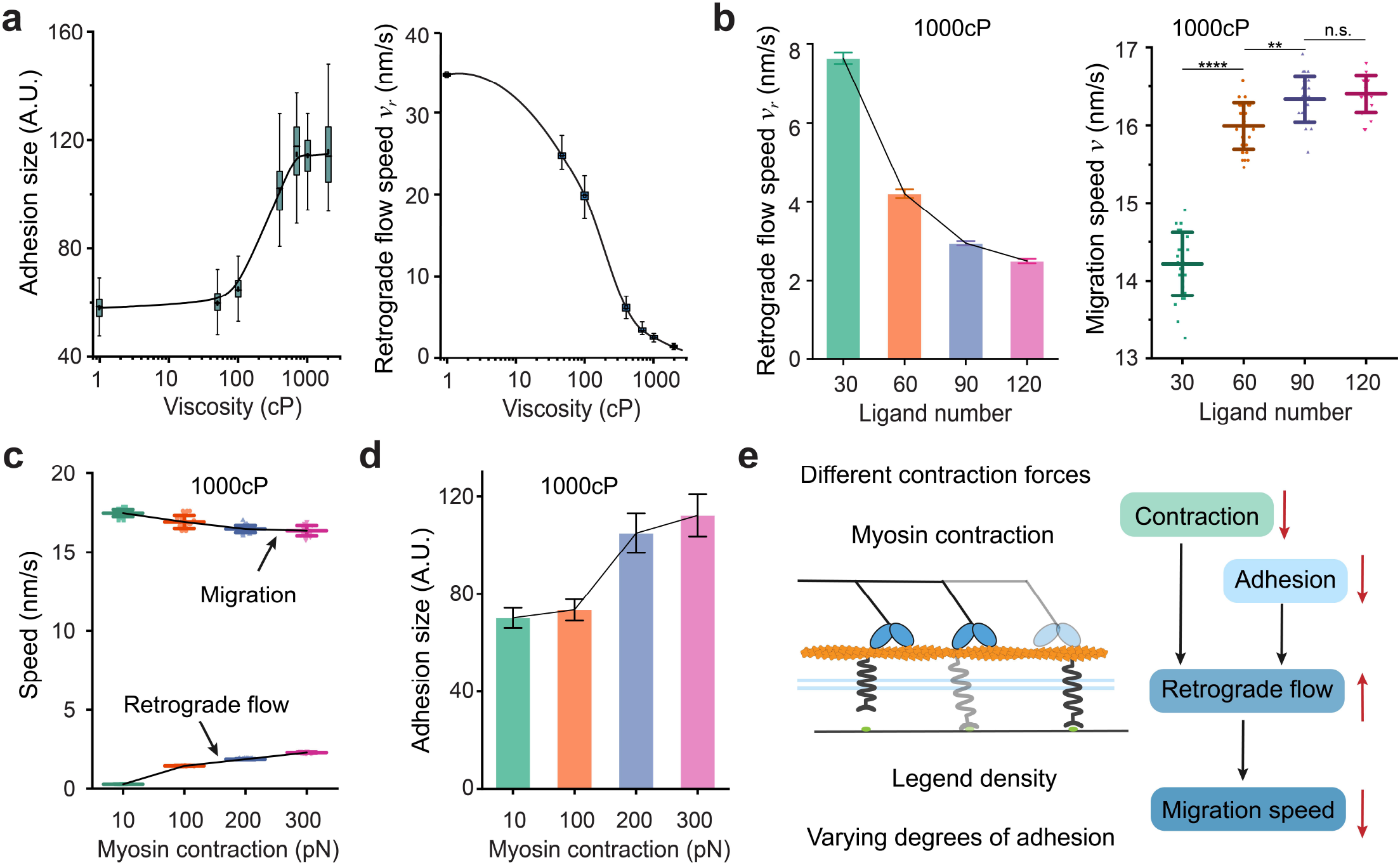
Mechanobiological cues for actin flow and cell migration. (a) Relation between ECE viscosity and cell-ECE adhesion (left) or retrograde flow speed (right). Whisker range 1 ∼ 99. (b) Relation between retrograde flow speed and the number of ligands for the high viscosity case (1000cP). Fewer ligands lead to fast actin flow (left), and correspondingly slower migration speed. When the ligand becomes dense enough, it has less influence on the migration speed. Error bars, s.d. (c) Effect of myosin contraction on retrograde flow and migration speed. Greater myosin contraction leads to slightly higher retrograde flow speed and slower migration speed. Error bars, s.d. (d) Effect of myosin contraction on adhesion. Within a certain range, increased myosin contraction leads to strengthened adhesion. (e) Schematic mechanism of ligand density and myosin contraction for actin flow and migration.

Myosin contractile forces can also affect retrograde flow and cell migration speed. In high viscosity ECM, increased myosin contraction leads to slightly faster retrograde flow (Fig. 5c), and this change in retrograde flow has a small effect on migration speed (Fig. 5c). That is, myosin contractility has little influence on migration speed in the high viscosity case, which is consistent with some experimental observations^22,48^. Besides, increased myosin contractility within a range can strengthen adhesion (Fig. 5d), because of the enhanced integrin binding mechanism^40^. As myosin contractility increases further, adhesion may become weaker (Fig. S7a). We summarize these relations with a schematic representation (Fig. 5e), illustrating how ligand density and myosin contraction collectively influence retrograde flow and migration speed. Higher ligand density enhances adhesions, slows retrograde flow, and promotes forward migration. On the other hand, elevated myosin contraction pulls on the AF network, accelerating retrograde flow and, as a result, slowing forward migration. While in the high viscosity ECM, the effect of myosin contractility has less effect because retrograde flow is very slow as adhesion strengthens, and AF polymerization is the main protrusion factor (Fig. S7b).

### Change in viscosity leads to mechanobiological migration memory

Cell migration memory is an emergent property that allows cells to utilize past experience to inform future movements, with significant impacts for pathological processes^24,25^.The cytoskeleton-based mechanical memory was reported in some experiments^49,50^, and might be related to the formation of long-term memory^51^. In this study, we elucidate the mechano-chemical coupled mechanisms of short-term cell migration memory. When a cell is transferred from the low viscosity ECE (1 cP) to high viscosity ECE (1000 cP), its migration speed initially drops sharply, indicating the immediate mechanical impact of the higher viscosity (Fig. 6a). As the cytoskeleton adapts by polymerizing and forming a denser AF network (Fig. 6c), the migration speed gradually increases. By this mechanism, the cell can regain its speed despite increased environmental resistance. Fig. 6b shows more distinctly that the cell moving in the low viscosity environment exhibits slow migration speeds, but it achieve faster movement through cytoskeletal adaptation when transferred to the high viscosity condition. This increase in migration speed under high viscosity conditions suggests that the cell requires sufficient propulsive force to overcome the added resistance. Our results show that the density of the AF network increases with rising viscosity (Fig. 6c), indicating that cells adapt to higher mechanical resistance by constructing a denser cytoskeletal structure, enabling them to exert stronger forces against the high viscosity ECE.

**Fig. 6.**
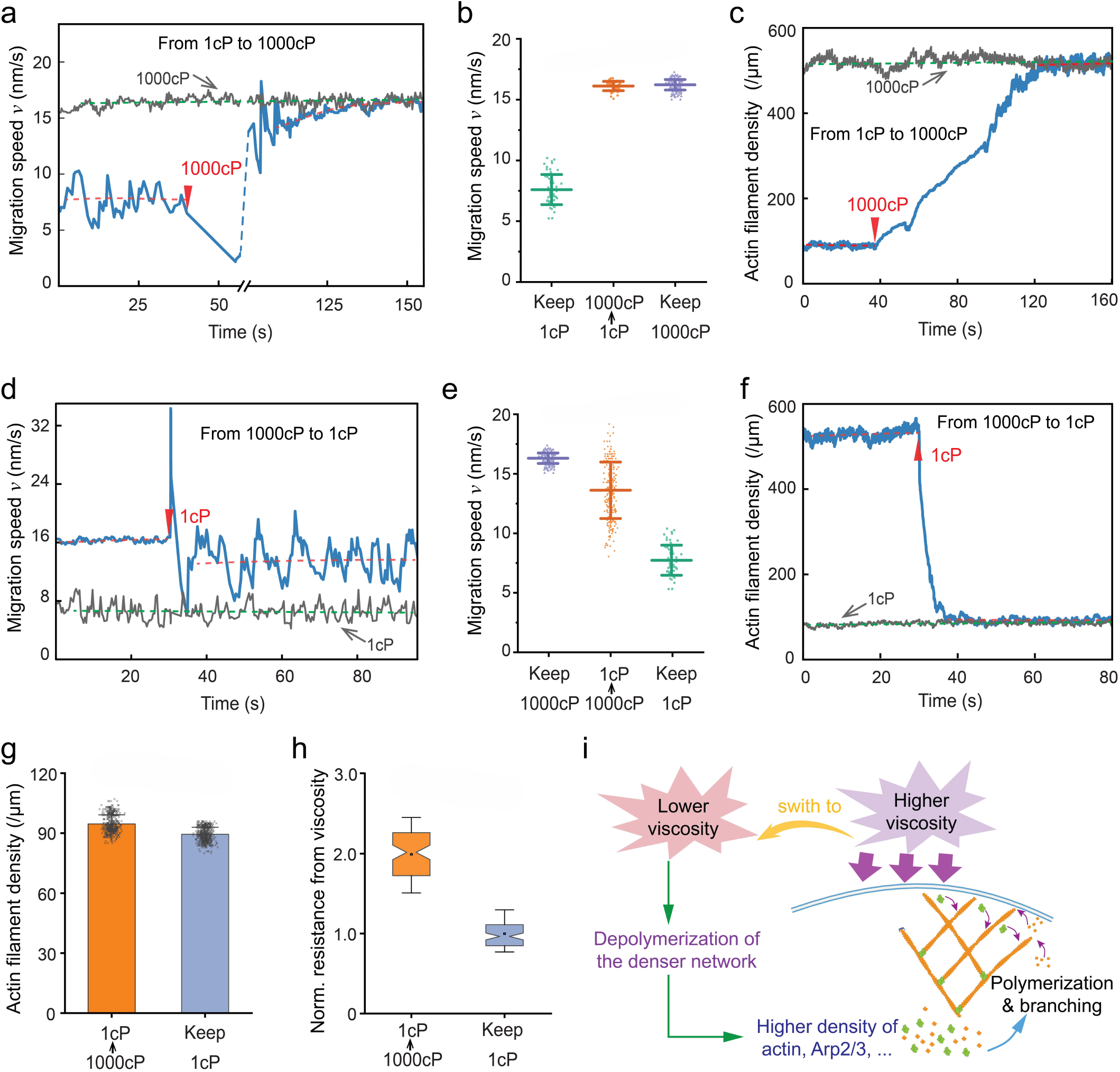
Changes in the viscosity of ECE and the effect on cell migration and cytoskeleton evolution. (a) The change in migration speed when a cell is transferred from low to high viscosity. (b) Comparison of migration speed in the viscosity changing and invariant cases. Error bars, s.d. (c) The change of AF density with viscosity. The AF density becomes larger with increasing viscosity. (d) The change in migration speed when a cell is transferred from high to low viscosity, with a sufficient actin monomer supply. (e) Comparison of migration speed in the viscosity changing and invariant cases. The migration speed retains some memory, that is, the cell maintains a similar speed in the high viscosity case after being transferred to the low viscosity case. Error bars, s.d. (f) The AF density changes as ECE becomes more viscous. (g) The difference in AF density for the case of transfer from high viscosity and invariant low viscosity. The small larger density is significant for the cell to maintain higher speed, which can provide greater propulsive force (h). (i) The short-term memory mechanism of cell migration. Cells in a high viscosity environment are stimulated to synthesize more relevant proteins, such as Arp2/3 and actin, to form a denser AF network. When a cell is transferred to a low viscosity environment, the cell still maintains the depolymerization of the dense AF network far from the leading edge, to produce a higher protein density, which is conducive to cytoskeletal polymerization. This in turn generates a greater propulsive force for cell movement. The time zero is calculated from the time after the Af network has produced stable propulsion. All traces in a, c, d and f are the trace after steady migration.

Interestingly, in the reverse case, when a cell is transferred from the high viscosity (1000 cP) to low viscosity (1 cP) condition, the migration speed first increases rapidly and then decreases. However, compared with the speed for cells consistently exposed to low viscosity, the migration rate stabilizes at a higher level, similar to that observed in the high viscosity condition (Fig. 6d). Fig. 6e compares the migration speeds in both constant and changing viscosity scenarios. The cell transitioning from high to low viscosity can maintain a higher migration speed than that under constant low viscosity conditions. This phenomenon suggests a short-term migration memory, wherein the cell can “remember” the motility developed in the high viscosity environment, allowing it to maintain faster movement. By interacting with epigenetic changes, this cytoskeleton-based memory may favor long-term memory formation^23^.

To further elucidate this memory effect, we examine the AF density when a cell is transferred from high to low viscosity. Although the AF density decreases upon transition to the low viscosity environment (Fig. 6f), it still remains higher than that of the cell always exposed to low viscosity conditions (Fig. 6g). Additionally, the cell transferred from high to low viscosity encounter greater resistance than that in continuously low viscosity environments due to their higher migration speed (Fig. 6h), which favors AF branching. This residual cytoskeletal density, coupled with elevated actin monomer concentration near the leading edge (Fig. S8), induces the increased migration speed in a low viscosity ECE. AF polymerization is influenced by actin monomer concentration near the leading edge, which is supplied from the cytosolic pool, and by recycling of depolymerized filaments (Fig. S8). Additionally, the enhanced contractility at the cell rear under high viscosity^23^ may further drive forward flow of actin monomer in the cytosolic pool to the leading edge^52^. In addition, the change and different fluctuations in actin monomer concentration can affect the duration of migration memory (Methods and Fig. S9). To explain this memory phenomenon, we propose a mechano-chemo-biological mechanism (Fig. 6i). When cells are exposed to a high viscosity environment, they increase the assembly of AF network that facilitates movement in resistant conditions. Even after transitioning to a lower-viscosity environment, this enhanced network assembly allows the cells to maintain a higher migration speed than those that were never exposed to high viscosity. This mechanical memory may be further supported by potential epigenetic changes within the cell (Fig. S10b), which requires further in-depth investigation.

## Discussion

Our model demonstrates a linkage between AF deformation and AF network formation through mechano-chemobiological mechanisms. Together with the consideration of retrograde flow, the model reveals a biphasic response in migration speed as ECE viscosity increases (Figs. 2a and S5). This model shows that increased viscosity can strengthen adhesions, reduce retrograde flow, and modulate actin polymerization dynamics, allowing the cells to sustain faster migration speeds up to a threshold where resistance becomes too high. This mechanistic insight underscores how adhesion strength and actin dynamics collectively mediate viscosity-dependent cell motility. The phenomenon, wherein cancer cells move faster in more viscous ECE, is consistent with experimental observations^20,22,23^. The biphasic pattern indicates that cytoskeletal adaptability is not only a function of the related proteins availability but is also tuned by the physical properties of the extracellular environment.

Actin polymerization is an essential process to generate propulsive forces for cell movement. The actin monomer availability is a key factor in driving AF polymerization. In low viscosity environments, cells require less actin polymerization to overcome low mechanical resistance, while in high viscosity environments, the demand for actin monomers increases as cells must exert greater force to migrate through a denser and more resistant medium. The efficient actin monomer recycling and transport ability^52,54–56^ helps maintain the local actin monomer concentration near the leading edge at a sufficient high level to gurantee continuous polymerization. Besides, elevated actin monomer concentrations promote rapid polymerization, translating to faster migration speeds (Fig. 4b). Biomechanical factors such as ligand density and myosin contractility also affect cell migration. Ligand density, which correlates with cell-ECE adhesion dynamics via integrins, can influence the adhesion strength, thereby impacting retrograde flow. Optimal ligand density facilitates effective adhesion, promoting motility. Myosin contractility also plays a central role in regulating retrograde flow. Increased myosin activity enhances cytoskeletal tension, driving retrograde flow. However, in high viscosity environments, retrograde flow is naturally slow, and the impact of myosin contractility can be diminished. This is verified by the experimental results that inhibition of myosin has a negligible effect on cell motility in high viscosity ECE^22^. This nuanced interaction between adhesion, contractility, and viscosity underscores the complicated mechanism, where cells regulate cytoskeletal tension and adhesion to achieve optimal migration.

Finally, this study reveals the mechanisms of the cytoskeleton-based short-term migration memory, wherein cells exposed to a high viscosity environment can maintain faster migration speeds even after transitioning to low viscosity conditions. This phenomenon suggests that cells can store information from their previous mechanical environment and use it to maintain enhanced motility. At the molecular level, this memory is likely related to the increased concentration of cytoskeletal proteins, such as actin monomers and Arp2/3 complexes, which is primarily due to rapid monomer transport and efficient actin recycling mechanisms (Figs. 6i and S8). The persistence of these proteins results in a denser AF network, providing sustained structural integrity and enabling the cell to move more quickly. A biological implication of this memory effect is particularly relevant in the context of cancer metastasis. During metastasis, cancer cells need navigate through diverse tissue environments, ranging from high viscous extracellular matrices to more fluid environments like blood vessels or lymphatics. Maintaining enhanced motility after transitioning from dense to softer tissues might be a significant ability of cancer cells, allowing them to more effectively disseminate and colonize distant sites. Our model demonstrates how biomechanical cues can drive short-term cytoskeletal memory after viscosity transitions, which is similar to the experimentally observed mechanical memory^49,50^. The biochemical pathway-medicated transcriptional changes^23,57^ may explain the long-term regulatory mechanisms for migration behavior. To fully integrate those mechanotransduction pathways and transcriptional feedback mechanisms^24,58^, future studies are needed to develop whole-cell models that encompass both cytoskeletal dynamics and transcriptional changes, including their interaction^51^. Such models would provide a unified understanding of how mechanical and biochemical cues synergistically regulate migration behavior across different time scales.

In summary, we have highlighted the effects of biomechanical cues, particularly ECE viscosity, on modulating cytoskeletal behavior, adhesion and cell motility, as shown in Table 1. We reveal how a cell dynamically integrates mechanical resistance with biochemical signals to tune its cytoskeletal structure and migration speed. Furthermore, the discovery of cytoskeleton-based migration memory introduces a new layer of complexity to our understanding of cellular adaptation, revealing how transient changes in the mechanical environment can induce effects on the migratory machinery. This work provides potential therapeutic insights for inhibiting abnormal cancer cell migration, suggesting that manipulating tissue mechanics or targeting the molecular pathways involved in AF polymerization and migration memory may offer novel strategies for limiting metastatic progression.

**Table 1.** Key cellular and subcellular behaviors and predictive capabilities of our model.

|  | Experimental phenomena | Our model |
| --- | --- | --- |
| AF Polymerization | Slower polymerization for larger resistance<sup>26</sup><br>Denser AF network for larger resistance<sup>26,29</sup> | Yes (Fig. S4)<br>Yes (Figs. 3c, S3) |
| Adhesion | Strengthened adhesion with increasing force applied to integrins<sup>40,53</sup><br>Increasing viscosity strengthens adhesion<sup>18,23</sup><br>Increasing viscosity slows retrograde flow<sup>22,23</sup> | Yes (Figs. 5d, S2)<br>Yes (Fig. 5a)<br>Yes (Fig. 5a) |
| Cell motility | Actin but not myosin dominates cell motility in the high viscosity condition<sup>22,48</sup><br>Increasing viscosity speeds up migration<sup>18,20,22,23</sup><br>Migration slows as viscosity becomes too high (not enough experiments) | Yes (Figs. 5c, S7b)<br>Yes (Figs. 2a, S5)<br>Yes (Figs. 2a, S5) |
| Memory effect | Cell migration speed remains high in low viscosity environment after pretreatment in high viscosity environment (long-term)<sup>23</sup><br>AF polymerization rate becomes faster in low resistance after pretreatment in high resistance condition (short-term)<sup>49</sup> | Long-term is not included<br>Yes (Figs. 6d, 6e, S9) |

## Methods

### Polymerization of the actin filament

AF polymerization is a dynamic process corresponding to the addition of actin monomers primarily at the barbed end and the removal of monomers at the pointed end. In the simulation, we assume that the process involves the addition of actin monomers only at the barbed end while the removal of actin monomers occurs at the pointed end. AFs polymerize over time and may be capped by capping proteins or interact with the membrane. The polymerization rate can be obtained from Eq. (3). The concentration of actin monomers *C*_*a*_ has an initial minimum value 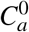. This concentration in the following simulation is related to the density of the AF network already far from the leading edge because the AF depolymerization can release actin monomers, which supplies free actin monomers^55,56,59,60^. Additionally, there is an upper limit to the concentration 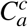 due to the diffusion limit of actin monomers. That is, the temporal concentration of actin monomers is determined by the AF density and the diffusion coefficient, corresponding to Φ and *γ* in Eq. (3), respectively. The force-dependent probability of AF polymerization is an exponential relation as in Eq. (S1). The association rate *k*_on_ and dissociation rate *k*_off_ of actin monomers to the polymerizing AF are constant during the simulation.

As the filaments polymerize, they might interact with the membrane. Branching may occur on the convex side of the mother filaments and new daughter filaments begin to polymerize at the Arp2/3 binding site^30,33^. The number of daughter filaments of a mother filament can be determined according to

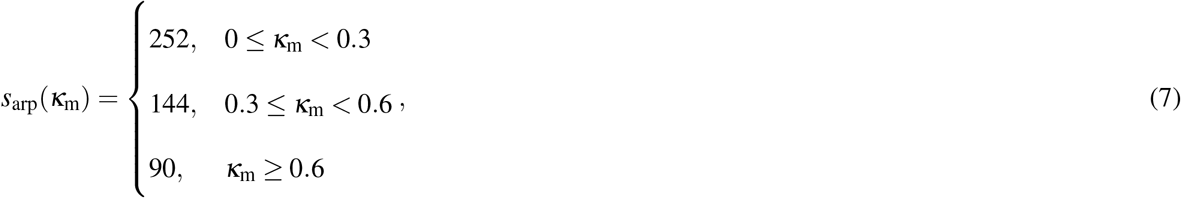

where *s*_arp_ is the distance between two Arp2/3 binding to a mother filament. Thus, the number of Arp2/3 binding to the filament is *N*_*arp*_ = ⌊*l/s*_arp_⌋, where ⌊·⌋ denotes the rounding down operator. Additionally, the critical curvature is set because the experimental observation shows that the bending angle is several degrees^34^. Besides, different types of cells might have different abilities to generate branched network, different *s*_arp_ can simulate these differences. Experimental results have shown that the angle between the branched daughter filament and mother filament is about 70^°7,61^. During the simulation, the branch angle is set as a random value between [68^°^, 72^°^].

### Retrograde flow

The polymerization of AF is also accompanied with the retrograde flow, which is simulated by the molecular clutch model in this study. The retrograde flow speed of the whole network is simply calculated using Eq. (6). The adhesion, corresponding to clutches in the molecular clutch model, is regulated by the ECE viscosity and the force acted on the clutch (Eq. (S14)). The adhesion strengthening is implemented by adding new clutches with a dynamic rate (Eq. (S10)). Then, the engaged clutch number is updated using Eq. (S8 - S10). Hence, the retrograde flow speed at the next time step can be updated using Eq. (6). The differences between the molecular clutch model used in this study and other developed related models are listed in Table S1, and the detailed formulation of the molecular clutch model can be found in the Supporting Information.

### Simulation process

At the start of the simulation, the starting points of the mother filaments are initialized. First, we select a 2-dimensional region with a width of *x* ∈ (−500, 500nm), assuming that the cell membrane is flat within this width range. The initial position of the cell membrane is *y* = 60nm. The polymerization starting points of the initial mother filaments are randomly generated and distributed in the height range *y* ∈ (0, 20nm) and the width range *x* ∈ (−500, 500nm). The number of initial mother filaments is *N*_mf_. To ensure that the number of mother filaments is roughly uniformly distributed over the width, the generation was done by dividing the width into 10 parts, and randomly generating *N*_mf_*/*10 coordinates of starting points in each of these parts. The orientation of AFs relative to the cell migration direction is generally around ±35^°62,63^. Thus, the initial angle of the generated mother filaments satisfies a Gaussian distribution *ϕ* ∼ *N*(±35^°^, 5^°^). Next, the AF polymerizes by adding individual actin monomers whose radius is *δ* = 2.5nm before growing to its maximum length or being capped, and the filament closer to the leading edge has a higher priority for assembly with actin monomer. The capping situation includes two scenarios, one is the randomness of generated length (a Gaussian distribution *N*(*l*, ±20nm)) and the other is the stopping of polymerization beyond a certain distance from the membrane. The polymerization rate is calculated using Eq. (3), where polymerization slows as actin monomer decreases. In the simulation, the initial polymerization rate is expressed as 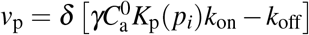. At each time step, the number of actin monomers is calculated by 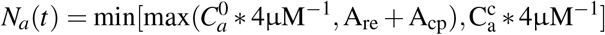, where *A*_re_ equals to the number of aged AFs divided by a random number ℛ (4, 6), i.e. the actin monomers from recycling. The pointed ends of the aged AFs are between 1000 nm and 1200 nm from the leading edge, i.e., actin filament depolymerization is assumed to occur one micron from the leading edge. For simplicity, the actin monomer from the cytosolic pool is set as *A*_cp_ = *A*_re_. When these actin mononers are depleted, the simulation proceeds to the next time step. Meanwhile, there is a retrograde flow whose speed is calculated by Eq. (6). The retrograde distance is Δ*s*_r_ = *v*_r_Δ*t* at each time step, where Δ*t* = 0.02 s is the incremental time. When the polymerizing AF contacts the membrane, it bends. There is an interaction force calculated by Eq. (5), which acts as the driving force for cell migration. When the propulsive forces of all AFs (left side of Eq. (1)) are greater than the resistance forces (right side of Eq. (1)), the cell migrates with a step size of Δ*d*. The migration speed is *v* = Δ*d/*(*t*_*j*_ − *t* _*j*−1_), where (*t* _*j*_ − *t* _*j*−1_) is the time from last to current migration step. Meanwhile, after bending of the AF, the branching phenomenon, that is, the binding of Arp2/3, will occur according to Eq. (7). Besides, the interaction force between the AF and the membrane will inhibit the polymerization speed in the next time step, as described by the function *K*_p_(*p*_*i*_). In summary, the flowchart of this simulation process can be found in Supporting Information Fig. S11.

To simulate the memory effect, the actin monomer concentration remains high (*C*_*a*_ ≃ 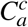 in Fig. 6 and different fluctuations in Fig. S9) near the leading edge after the cell has transferred from the high to the low viscosity condition, based on the mechanism shown in Fig. S8. The larger resistance in the high viscosity ECE leads to denser F-actin networks to push the membrane forward^26,29^. The branched Arp2/3–actin filament network subsequently leaves the ‘activation zone’ as the membrane is pushed forward. Then the aged AFs undergo debranching and depolymerization, and the actin monomers released by the AF network disassembly can be reused for subsequent rounds of polymerization^55,56,59,60^. The actin monomer transportation can also support continuous polymerization^52,54^.

## Acknowledgements

Supports from the National Natural Science Foundation of China (Grant Nos. 12032014 and T2488101) are acknowledged. Z. Lin hopes to thank Dr. Shuang Li and Dr. Wenyu Kong of the Tsinghua University for their helpful discussions.

## Author contributions statement

Z.L., X.C. and X.-Q.F. designed research; Z.L., X.C. and X.-Q.F. developed computational framework; and Z.L., X.C. and X.-Q.F. analyzed data and wrote the paper.

